# Network analysis of lymphocyte nucleus staining image —Data mining of lymphocyte image

**DOI:** 10.1101/396143

**Authors:** Da-Dong Li, Xing-Lin Yang, Qian-Yu Xiong, Yue-Dong Liang, Shui-Qing Liu, Hai-Yan Hu, Xiang-hong Zhou, Hai Huang

## Abstract

**Background**: A complex network has been studied and applied in various disciplines. As network analysis and image processing are based on matrices, this research analysed the changes in the chromatin image of lymphocyte nuclei in peripheral blood of humans using a network motif and static features (static parameters), so as to complete image classification with network method.

**Methods**: Image processing technology was used to establish a chromatin image network of a cell nucleus; Network analysis tool Pajek was used to display the special motif of an isolated structural hole with different symmetric line values; afterwards, the frequency of occurrence of this structural hole in patients with nasopharyngeal carcinoma and AIDS, and healthy people was computed. Then by applying the network static features as variables, the chromatin images of stained lymphocytes from the three groups of people were classified and recognised by using an extreme learning machine (ELM).

**Results**: The frequency of occurrence of the isolated structural hole with different symmetric line values was adopted to distinguish the structures of the chromatins of peripheral blood lymphocytes in patients with nasopharyngeal carcinoma and AIDS, and healthy people. Similarly, The static features of the chromatin image network of a cell nucleus were applied to classify and recognise the morphological and structural changes in chromatins for peripheral blood lymphocytes in the three groups of people.

**Conclusion**: The surface chemical and physical characteristics, as well as the polymerisation link status of biomacromolecules such as DNA, RNA, and protein in the lymphocyte nucleus change under certain pathological conditions. The change influences the combination of small molecular staining materials and any associated biomacromolecules. Therefore, various macroscopic and microscopic changes were found in the chromatin images of the cell nucleus. The microscopic changes include the variations of the extent of staining of chromatin in the nuclei, coarseness and direction of the texture therein, the size of stained conglomerations, *etc*. These changes contribute to the differences in chromatin image networks among the same type of cells across the three groups. Based on this, the model can be used to classify and reorganise certain diseases. The results prove that using complex network to analyse the chromatin structure of a cell nucleus is of significance.

## 1. Introduction

Network science, as a new interdisciplinary subject, provides a unified framework for the recognition and processing of complex systems [1]. Concerning the application of network characteristics to the explanation of biological, physical, and social phenomena, scholars have constructed analytical models and described these phenomena using the static or dynamic features of their networks. Network analysis technology, based on graph theory, has become an important tool in the investigation of complex systems including sociology, physics, and cell biology [2]. The application of network models in the interpretation of biological complex systems is now increasing [3].Cells, as an integral unit in living systems, can be regarded as a complex system. The authors consider that when blood cells are chemically stained using chemical dyes such as Wright’s stain, the staining materials can be regarded as an extrinsic probe composed of small molecules in chemico-biology [4,5]. Due to the combination of staining molecules with the biomacromolecules in the cell, the structural information including the extent of staining of the chromatin, the texture, and size of stained conglomerations of the stained cells can be observed by microscopy. Therefore, based on image processing technology, a weighted adjacent matrix which reflected the bonding strength of the pixel was converted from the chromatin images of a stained cell nucleus. In this way, an undirected biological network, called the chromatin image network of a cell nucleus, and the graphical network model which could be analysed using mathematical language were constructed. By analysis using network technology, the authors found some special structural holes (three-node motifs whose two nodes were not connected.It’s a triangle without an edge) within the network. As these structural holes were not connected to any other nodes in the network, they were called isolated structural holes(visually, this article refers to a triangle with one edge missing) (see Figure 3,4). Meanwhile, the values of the two lines in the structural holes were flexible. The structural holes were mainly found in the medium and small lymphocytes in patients with AIDS and nasopharyngeal carcinoma. Therefore, the chromatin image network of the lymphocyte nucleus of these patients significantly differed from that in healthy people. Afterwards, some static features of the chromatin image network of the cell nucleus were calculated and it was proved using an extreme learning machine (ELM) [6,7,8] that these features were effective in the classification and recognition of chromatin images of lymphocyte nuclei of patients with AIDS and nasopharyngeal carcinoma, and healthy people.

## 2. Materials and methods

### 2.1 Materials

Wright’s chemical stains were used. Blood smears were prepared using a fixed quantity of peripheral blood.

### 2.2 Methods

#### 2.2.1 Staining of cells and acquisition of cell image samples

In the preparation of blood smears in the laboratory, 10 μl of blood was used each time. After the blood smears were dried, drops of staining solution and buffer were added to stain the blood cells. After 9 min of staining, the blood smears were washed using distilled water and then dried in the air for spare use. The staining solution and buffer used were produced in the same batch.

The images of the isolated peripheral blood lymphocytes in the blood smears were taken using an OLYMPUS CX41 microscope equipped with a high-resolution digital micro-camera. An oil lens was used in the acquisition of each image. In this process, isolated single nucleus cells needed to be found first. After confirming that the cell was a lymphocyte by its morphology, the cell was placed to the centre of the view-finder before being photographed. The intensity of the light source on the camera was constant throughout. The acquired cell images are shown in Figure 1. At least 10 images of the medium and small lymphocytes in the peripheral blood were taken for each sample (10 images).

**Figure 1.**
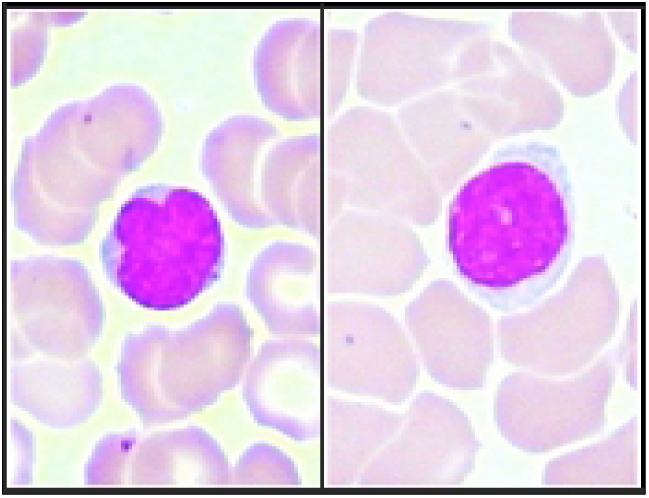
Peripheral blood lymphocytestaken in two different views.

#### 2.2.2 Construction of the chromatin image network of lymphocyte nucleus

The images were trimmed to show single nucleus cells (see Figure 2 A).Afterwards, chromatin images (measuring 50 × 50 pixels) were taken from the trimmed images of single cells. They were obtained thus: the cell nucleus images were separated from the cell images in Figure 2 using the modified *k*-mean clustering method; the centroids of the cell nuclei were found and used as central points to trim images to 50 × 50 pixels. Repeated experiments revealed that when the trimmed area was larger than 50 × 50, the area was likely to cover the image of the cytoplasm for small cell nuclei. This kind of trim was invalid. The process is shown in Figure 2B, in which, the red point in the gray cell nucleus is the calculated centroid of the image and the red dashed box represents the trimmed area. The last image in Figure 2B was the 50 × 50 pixel version of the trimmed chromatin image with the centroid aligned in the middle; it was then named and in .jpg format stored.

**Figure 2.**
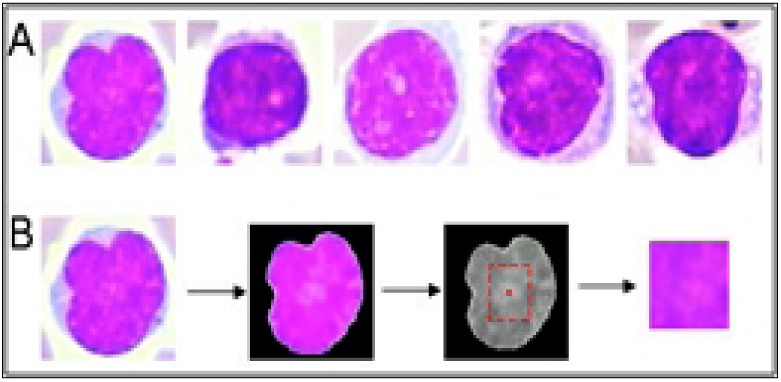
(A) Trimmed images of five single cells;(B) Process for acquiring 50 × 50 pixelchromatin image of lymphocyte nuclei.

#### 2.2.3 The display of a special isolated structural hole(Extraction of isolated structural holes from the chromatin image network of a cell nucleus)

In the process of converting the gray-scale image of the cell nuclear chromatin into the adjacency matrix of the network, we find an optimal threshold to generate the adjacency matrix of the network, so we find that the adjacency matrix can show some isolated structural holes in the chromatin image network of a cell nucleus were displayed using Pajek.

As mentioned, isolated structural hole here meant that the motifs were independent and were not connected to other nodes in the network. They were small-scale homomorphic sub-graphs.

The process was as follows:

Step 1: The 50 × 50 chromatin images were converted to gray-scale images. Adaptive filtering was applied.

Step 2: Complete the image intensity (gray) was adjusted, find the optimal thresholds,look-up the main pixels, conversion to binary image, formation the adjacent matrix of the chromatin image of a cell nucleus,the adjacent matrix was converted to a .*net* document which was recognisable by Pajek.

Step 3: After locating the .*net* document in the relevant folder, Pajek 2.00 (November 1996 - October 2010) was started. After unfolding the folder containing the .*net* documents of the adjacent matrices of the chromatin image networks by clicking File/Network/Read in the menu bar of Pajek, the .*net* document for analysis was selected. Then it was read in Pajek by clicking the Open button in the dialog box.

Step4: The Pajek graphic window popped-up by successively clicking the Net/Partitions/Core/All, Partition/Make Vector, and Draw/Draw-Partition-Vector in the menu bar. Then by clicking the Layout/Energy/Kamada-Kawai/Separate Components (shortcut key: Ctrl+k) in the menu bar of the graphics window, a network of the chromatin image of a nucleus was displayed. The result, which was stored by clicking File/Pajek Project File/Save in the menu bar, could be accessed by clicking File/Pajek Project File/Read thereafter. To output, and store, the images, the Export/2D command in the graphic window of Pajek could be run. The command provides three formats (including Bitmap) for output and storage (see Figure 3).

**Figure 3.**
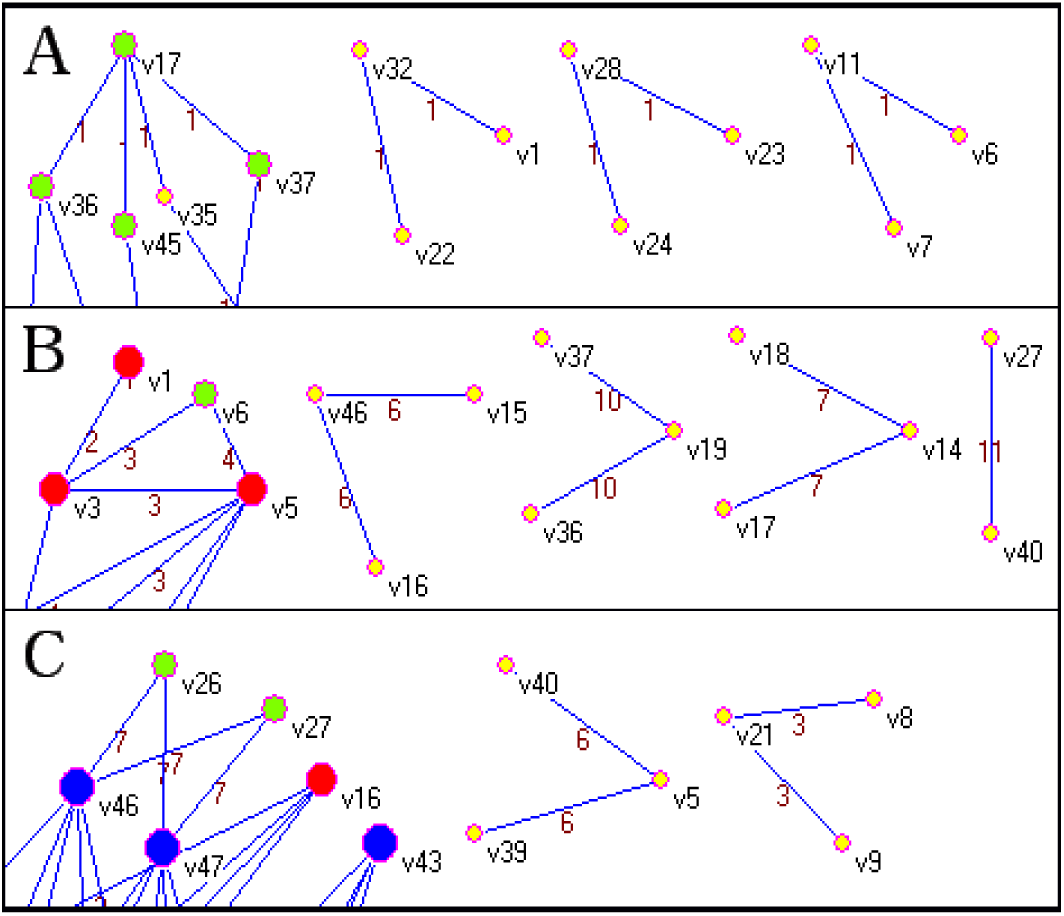
(A)Three isolated motifs (structural holes) in the network,(B) Isolated structural holes with three different symmetric line values,(C) Isolated structural holes with two different symmetric line values.

**Figure 4.**
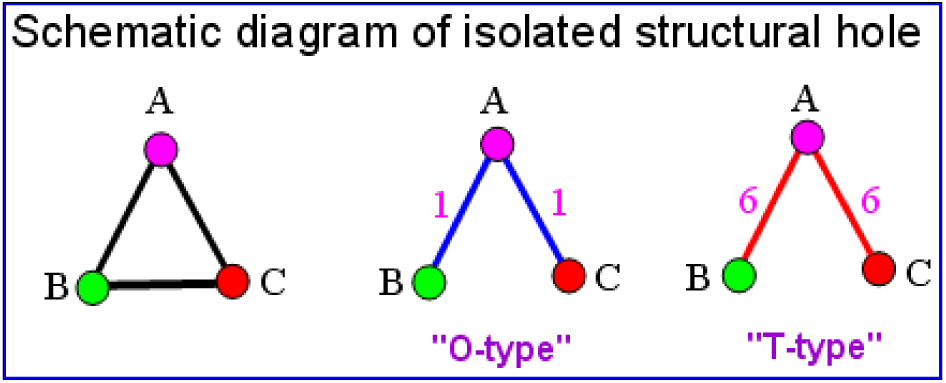
Motif and structural hole composed of three.

It’s very obvious. Figure 3A shows that there were three isolated structural holes in the network and that their nodes were v1-v32-v22, v23-v28-v24, and v6-v11-v7, respectively. Attention, that the symmetric line values of the isolated structural holes in Figure 3A are equal to 1, the symmetric line values of the solitary structural holes in Figure 3B and Figure 3C are all greater than 1,for example, in Figure 3C the symmetry line value of the isolated structural hole (v40-v5-v39) is equal to 6. In order to facilitate analysis of the chromatin image network of the lymphocyte nucleus, we temporarily refer to isolated structural holes with symmetry line values equal to 1 as “O-type”, Symmetric line values greater than 1 are called “T-type”.

#### 2.2.4 Statistics of frequencies(probabilities) and line values of isolated structural holes in different types of network

Samples: 30 healthy people,30 patients with clinically diagnosed nasopharyngeal carcinoma, 30 patients with clinically diagnosed AIDS. The results showed that the chromatin image networks of the three groups differed significantly (see Table 2 and Figure 5).

**Figure 5.**
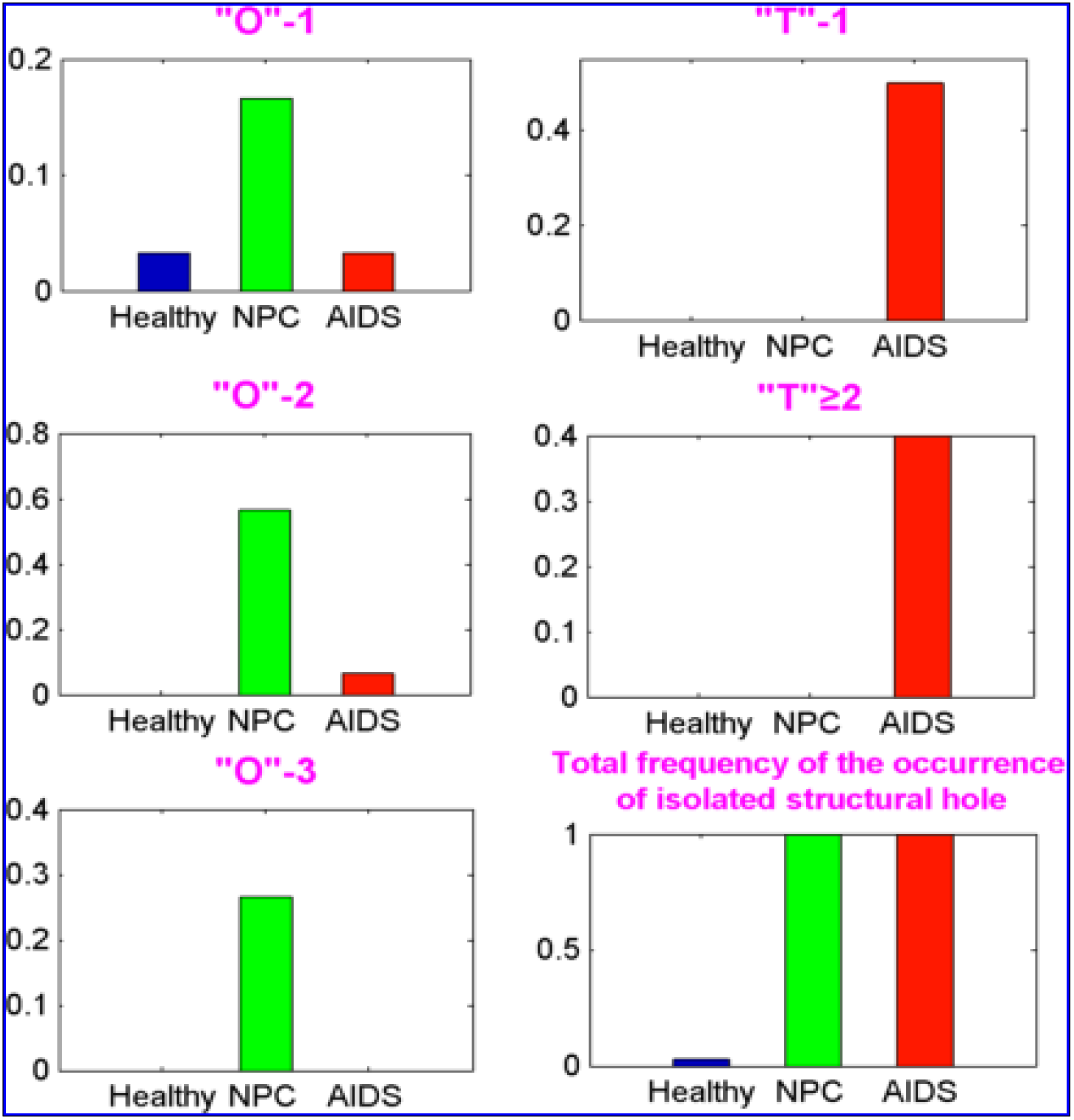
Frequency bar graph.

#### 2.2.5 The static features of the chromatin image network of nuclei were used to classify the images

Section 2.2.4 above counted the frequencies of “O” and “T” types of special motifs in the chromatin image network of the lymphocyte nucleus of healthy people, AIDS patients and nasopharyngeal carcinoma patients. The following is the classification of the chromatin image of the lymphocyte nucleus using the static features of the network. First step, use the adjacent matrix of the chromatin image network of nuclei to calculate the static features of some networks. In this paper, 23 network static features such as the average path length of the network are calculated. These static features were adopted as independent variables for classification. Second step, the back-propagation (BP) neural network is used to complete the variable screened for 23 network static features. The third step is to complete the classification of the chromatin image of the lymphocyte nucleus with the extreme learning machine (ELM) as follows:

In this study, we calculated a total of 23 network static features. This is to ensure that there are sufficient data and the diversity of data to satisfy the requirements of variable screening. These 23 static features of the network were loaded into the BP neural network for variable screening. Finally, 4 network static features were selected for image classification (these 4 network static features will be introduced later).

The number of lines in the network, which was computed using Pajek, was applied as the dependent variable, while the 23 static features were independent. Then the mean impact value (MIV) [9, 10,11] exported by the BP neural network was used to screen the independent variables. Based on MIV, a positive or negative value was generated for each independent variable (see Table 1).

**Table 1.**
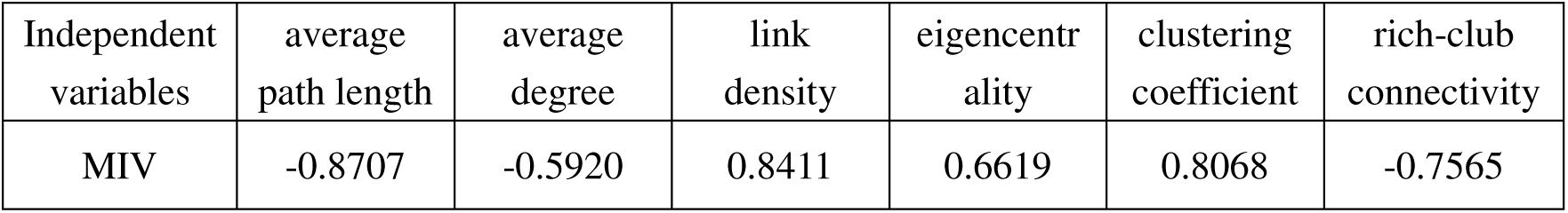
Six independent variables (static network features) with largest absolute MIV valuesTable 1 lists six independent variables with the largest absolute values of MIV screened from the 23 static features. The data revealed the influence, and its significance, of each independent variable on the values output by the BP neural network: independent variables with larger absolute values of MIV exerted larger influences on the output, while those with smaller absolute values of MIV showed less influence. Therefore, static features with large absolute values of MIV were selected as variables to recognise different types of chromatin images of nuclei using ELM. In this research, 30 patients with AIDS, 30 patients with nasopharyngeal carcinoma, and 30 healthy people were studied. That is to say, there were three types of data for classification and recognition and 90 data points for each static feature (independent variable). In the classification and recognition process, the static features were added step-by-step. Firstly, two static features (average path length and link_density) with the largest absolute values were selected to construct the data set for classification, that is, a data matrix measuring (90 × 2). A prediction accuracy was obtained at each time-step in the ELM operation, and a total of 10 times were used to compute the average accuracy. Then another static feature was added to establish a data matrix measuring (90 × 3). Likewise, the operation was carried out 10 times using the ELM to calculate the average accuracy. This process was and an independent variable was added continuously until the average accuracy was reduced. In this research, the computation finished as a data set measuring (90 × 6) was obtained, and therefore a data classification and recognition matrix composed of six independent variables was constructed. To ensure the objectivity and verisimilitude of the experiment, the test set and training set in each experiment were generated randomly. Thereby, nine sample test sets and 81 sample training sets were randomly produced to classify and recognise the three types of chromatin images.

| Independent variables | average path length | average degree | link density | eigencentality | clustering coefficient | rich-club connectivity |
| --- | --- | --- | --- | --- | --- | --- |
| MIV | -0.8707 | -0.5920 | 0.8411 | 0.6619 | 0.8068 | -0.7565 |

**Table 2.** Frequencies and line values of isolated structural holes in different types of network Where “O”-1 indicates that one “O” type isolated structural hole was detected, “O”-2 indicates that two “O” type isolated structural holes were detected, and “O”-3 indicates that three “O” types were detected. “T”-1 indicates that one “T” type isolated structural hole is detected, and “T”≥2 indicates that two or more “T” type isolated structural holes are detected.

|  | Frequency of one isolated structural hole (symmetric line value = 1) in the network ("O"-1) | Frequency of two isolated structural holes (symmetric line value = 1) in the network ("O"-2) | Frequency of three isolates structural holes (symmetric line value = 1) in the network ("O"-3) | Frequency of occurrence of an isolated structural hole (symmetric line value > 1) in the network |  | Total frequency of the occurrence of isolated structural hole |
| --- | --- | --- | --- | --- | --- | --- |
|  |  |  |  | 1 ("T"-1) | ≥ 2 ("T" ≥ 2) |  |
| 30 healthy people | 1/30=0.033 | 0 | 0 | 0 | 0 | 0.033 |
| 30 patients with NPC | 5/30=0.166 | 17/30=0.566 | 8/30=0.266 | 0 | 0 | 0.998 |
| 30 patients with AIDS | 1/30=0.033 | 2/30=0.066 | 0 | 15/30=0.5 | 12/30=0.4 | 0.999 |

Experimental results indicated that optimal classification and recognition effect was obtained using the data matrix composed of four static features (independent variables), including average path length, link_density, clustering coefficient, and rich-club connectivity (see Table 3). The definitions of, and computation methods for, the four features are described as follows:

**Table 3.**
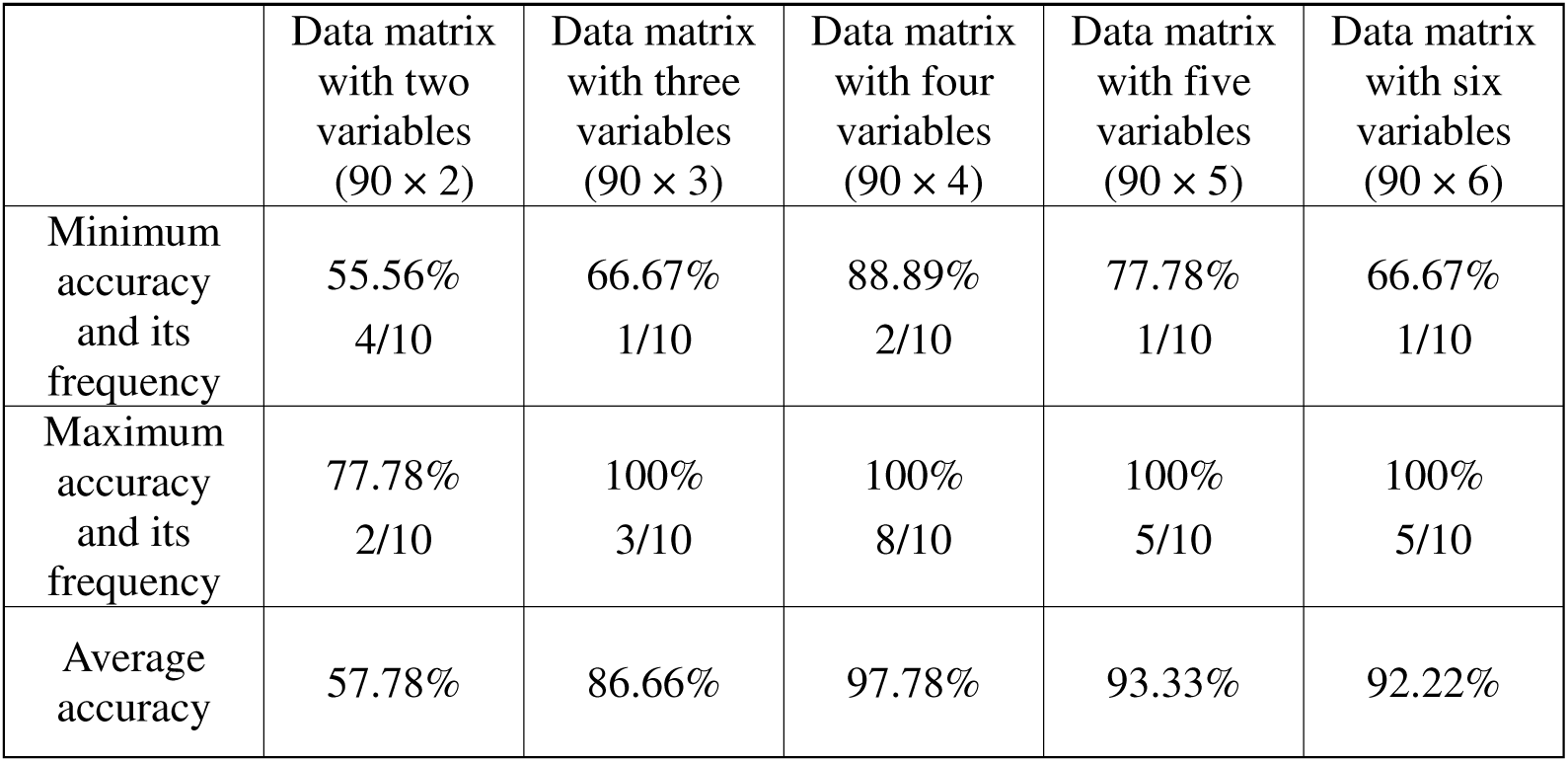
Classification and recognition results based on data matrices with different numbers of variables Table 3 indicates that the data matrix composed by four static features showed optimal classification and recognition effect, with a minimum accuracy of 88.89% and maximum accuracy of 100%. The minimum and maximum accuracies occurred two and eight times in 10 trials, respectively. It verified that the static features of the chromatin image network of cell nuclei can be applied to classify and recognise the morphological and structural changes in the chromatin of certain peripheral blood lymphocytes.

1.Average path length (*L*): Suppose that the shortest path length between any two nodes is *L* is*D*_*ij*_, then the average path length

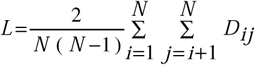

Where *N* is the number of nodes, the average path length refers to the average distance of all node pairs and reflects the average extent of separation of the nodes in the network. The shorter the average path length, the faster the transformation of information between nodes. The average path length is an important property of networks[12,13].

2.Link-density (*d*): It is the ratio of the actual number of lines (*m*) in the network to the maximum lines possibly contained in the network, that is, *d* = 2*m*/*N*/(*N—*1), where *N* is the number of nodes. Link-density represents the tightness of nodes in the network. The more lines in the network, the denser the network. A commnnity usually has relatively high link-density in social network[14, 15,16].

3.Clustering coefficient(C): The clustering of a complex network consists of triangular links, that is, a clustering coefficient is based on triple nodes. A triple has two kinds of structures: one is the closed triple, namely a complete triangle, constructed by three undirected lines connecting three nodes, and the other is an open triple (structural hole) whose three nodes are connected by only two lines. The clustering coefficient here means the number of closed triples in all the triples (including open and closed triples) and is defined as:

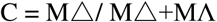

Where M△and MΛ represent the number of closed triples and open triples, respectively. The clustering coefficient reflects the collectivisation of the network. The larger the clustering coefficient, the greater the collectivisation of the network. The clustering coefficient is used to measure the transmitting capability of the network for local information[17,18,19].

4.Rich-club connectivity(Rich-club coefficient) (Φ (r)): there are several nodes with a large degree of connectivity (connecting with large amount of lines) in the network. These are rich nodes and have a high probability of being connected to other rich nodes. Therefore, these rich nodes constitute a rich-club. The connections among the rich nodes are described using rich-club connectivity, that is,

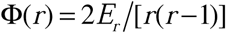

Φ (r) defines the ratio of the number (*E*_*r*_) of actual lines among the richest nodes in the number of *r* to the possible number *r*(*r* - 1)/2 of lines among these nodes in the network [20,21,22,23].

Here, it is necessary to introduce a brain network: similar to establishing a network using image processing technology, a brain network constructs a network using non-invasive imaging technologies such as magnetic resonance imaging [24,25]. The rich-club has been extensively studied and applied in research into brain networks. For example, it has discovered that rich-club hubs play an important role in information exchange in the human brain, while the link density of rich-club hubs of schizophrenia patients is diminished significantly [26,27,28].

## 3. Results

After years of sample collection, continuous exploration, and modification of the computer-aided numerical experimental method, the authors analysed the chromatin image networks of the peripheral blood lymphocytes of 30 patients with clinically diagnosed nasopharyngeal carcinoma, 30 patients with clinically diagnosed AIDS, and 30 healthy people. The results showed that the chromatin image networks of the three groups differed significantly.

### 3.1 The isolated structural hole in the chromatin image network of lymphocyte nucleus

The network motif was proposed by Israeli scientist Alon *et al*[29]. Motifs exist in all kinds of complex networks, such as biochemical networks, evolutionary networks, engineering networks, *etc*[30]. Therefore, discovering and recognising both motif and module, with their certain characteristics, in a network and investigating their characteristics, behaviour, and functions is important[31,32,33,34,35,36]. A motif which comprising three nodes A, B, and C is shown in Figure 4. The three nodes are connected to each other with three lines and form a triple in a social network. In the regulatory network of gene transcription, this kind of motif is also found, including feedforward, and feedback, loops [37]. This motif is also applied in research into cancer, *etc*[38,39,40]. As illustrated in the right-hand part of Figure 4, if nodes B and C are not connected, it seemed that there was a hole in the network. Therefore, the structure was called a structural hole. The structural hole concept was proposed by American sociologist Ronald Burt [41] and has been widely applied in sociology [42,43,44]. As B and C are not connected, there is no information exchange between them. Based on the method presented in Section 2.2.3, the structural hole was independently displayed as an isolated structural hole. The judgement standard for isolated structural holes is demonstrated in Figure 4. Firstly, it is independent, that is, it is not linked to any other nodes in the network. Secondly, it is composed of three nodes, among which two nodes are not connected. Thirdly, the three nodes show the same degree of connectivity. It was also observed from the chromatin image network of cell nuclei that the line values of the two lines in the isolated structural hole vary depending upon prevailing conditions. For example, the two lines v11-v6 and v11-v7 of the isolated structural hole with nodes v6-v11-v7 in Figure 3A were symmetrical and coincident: both the line values were unity. While, there were also isolated structural holes with different symmetric line values of 3, 4, 5, 6, 7, and 10, as demonstrated in Figures 3B and 3C. In Figure 3B, the isolated structural hole had three different symmetric line values (6, 10, and 7), and Figure 3Cshows isolated structural hole with two symmetric line values of 6 and 3. The frequencies and line values of the isolated structural holes existing in the chromatin image networks of healthy people, patients with nasopharyngeal carcinoma, and with AIDS were calculated (see Table 2 and Figure 5).

Isolated structural holes were not detected in all of the chromatin image of the lymphocytes of thepatients. Generally, among the 10 medium and small lymphocytes of patients with nasopharyngeal carcinoma and AIDS, isolated structural holes were detected in the chromatin image of one or two lymphocytes. Mostly, they were detected in the chromatin image of a single lymphocyte from each patient.Therefore, in the 10 chromatin image of the lymphocytes, the first image that detected an isolated structural hole was used as an analysis sample(First discovery principle). For healthy people, if the 10 chromatin image of the lymphocytes cannot detect isolated structural holes, the last 10th chromatin image of the lymphocytes is used as an analysis sample.

According to the results, the authors believed that the frequency of occurrence of isolated structural holes with different symmetric line values can be applied to classify, and describe, the chromatin structures of certain peripheral blood lymphocytes of patients with AIDS, nasopharyngeal carcinoma, and healthy people.

### 3.2 Classification and recognition of the chromatin images of cell nuclei of patients with AIDS, nasopharyngeal carcinoma, and healthy people using ELM

The method for constructing data matrices by adding static features (independent variables) step-by-step was described in Section 2.2.5. When a data matrix was built, the ELM was operated some 10 times. The classification and recognition results based on the data matrices with 2, 3, 4, 5, and 6 variables are summarised in Table 3.

## 4. Discussion

The research is still in its initial stages, and to some extent, this is a working report. As the applied samples were cells collected from patients with clinically diagnosed AIDS and nasopharyngeal carcinoma, whether or not the isolated structural holes were caused by medical treatment or other reasons was not revealed.

The statistical results in Table 2 and Fig.5 demonstrate that the isolated structural holes are effectual in the analysis of the changes in the chromatin of the lymphocyte nucleus. However, since there are only 10 nuclear chromatin images per case, and the first discovery principle is used in the detection, these will result in false negative (i.e. “T” and “O” -2, “O”-3 can not be detected). Therefore, it can be said that: 2.2.5 The static features of the chromatin image network of nuclei were used to classify the images, that is, to use the image classification method of the static characteristics of the network as a supplement and complete to the analysis of the isolated structural holes. Therefore, in order to further improve the analysis efficiency of isolated structural holes, it is necessary to increase the sample size and improve the detection method. In addition, it is too simple to divide the isolated structural holes into two types: “O” and “T”, which need further study and improvement. Besides, There is the sample of “healthy people”, which is only the result of screening of some physical examination items. That is, “healthy people” are not necessarily completelyhealthy.

It has been proved that a complex network is effective in analysing the chromatin images of peripheral blood cells. In particular, it has been verified that DNA is likely to be harmed by biological, physical, and chemical factors. As early as the 1960s, it has been found that viral infection shows a cytogenetic effect in cells of patients with viral diseases and human cells cultured in vitro. In other words, viral infection induces aberrations in human chromosomes [45,46,47]. Alkaline single cell microgel electrophoresis assays for eukaryotic cells, proposed by Singh *et al*., also known as a comet assay, is used to detect damage in the DNA of eukaryotic cells[48,49]. It is also a universal method for measuring multiple occurrences of damage in DNA and has been widely used [50,51,52]. Based on this, the authors believed that the occurrence of isolated structural hole in the chromatin image networks of certain lymphocyte nuclei, the differences in line values, and the variation of the static features of the network also reflected the fact that the DNA in the lymphocyte nuclei was damaged by certain pathological factors. It can also be explained by certain changes that occur to the higher structure, degree of clustering, linkage, and surface chemical and physical characteristics of macromolecules such as DNA, RNA, and those proteins in the lymphocyte nucleus changing under certain pathological conditions. These changes influenced the combination of staining materials’ small molecules and macromolecules and then caused the changes in the chromatin images (such as coarseness and direction of texture and the size of stained conglomerations) of the nucleus. Therefore, the chromatin image network changes and differs from that of the lymphocytes of healthy people. Furthermore, because macromolecules such as DNA show certain spatial structure in their cell nuclei, the spatial structure of these macromolecules can be observed in the images demonstrating the combination of small molecule staining materials and macromolecules. By using complex network technology, the network status can be analysed to obtain spatial information about the macromolecules. So, using complex networks to analyse the chromatin structure of cell nuclei was significant.

